# Pooled long-read sequencing for structural variant characterization in schistosome populations

**DOI:** 10.1101/2025.01.25.634855

**Authors:** Shalini Nair, Xue Li, Timothy J. C. Anderson, Roy N. Platt

**Affiliations:** Texas Biomedical Research Institute, San Antonio, Texas, USA

**Keywords:** *Schistosoma mansoni*, Nanopore, population genetics, parasite variation, quantitative trait loci

## Abstract

Pooled sequencing provides a rapid cost-effective approach to assess genetic variation segregating within populations of organisms. However, such studies are typically limited to single nucleotide variants and small indels (≤ 50bp), and have not been used for structural variants (SVs; >50bp) which impact large portions of most genomes and may significantly impact phenotype. Here, we examined SVs circulating in five laboratory populations of the human parasite *Schistosoma mansoni* by generating long-read sequences from pools of worms (92 -152 per population). We were able identify and genotype 17,446 SVs, representing 6.5% of the genome despite challenges in identifying low frequency variants. SVs included deletions (n=8,525), duplications (n=131), insertions (n=8,410), inversions (n=311), and translocations (n=69) and were enriched in repeat regions. More than half (59%) of the SVs were shared between ≥4 populations, but 12% were found in only one of the five populations. Within this subset, we identified 168 population-specific SVs that were at-or-near fixation (>95% alternate allele frequency) in one population but missing (<5%) in the other four populations. Five of these variants impact the coding sequence of 6 genes. We also identified 8 SVs with extreme allele frequency differences between populations within quantitative trait loci for biomedically important pathogen phenotypes (drug resistance, larval stage production) identified in prior genetic mapping studies. These results demonstrate that long-read sequence data from pooled individuals is a viable method to quickly catalogue SVs circulating within populations. Furthermore, some of these variants may be responsible for, or linked to, regions experiencing, population-specific directional selection.

**Significance Statement:** Structural variants (SVs) are large genomic variants that are frequently overlooked despite being the largest source of genetic variation within a population. This is because large SVs are expensive and difficult to genotype relative to single nucleotide variants or small indels, so are typically overlooked in population studies. This study attempts to solve these problems by using pooled samples and long-read sequencing to survey SVs circulating in five laboratory populations of the human parasite, *Schistosoma mansoni*. We were able to identify 17,446 SVs that impact 6.5% of the genome. A number of these SVs may be linked to population-specific adaptations. We also found 8 SVs that were associated with known parasitic traits from previous studies. This work highlights the value of long-read sequencing of pooled samples to document genetic diversity and provides a new method for exploring the role of SVs in parasite evolution and pathogenicity.

## Introduction

Structural variants (SVs), including insertions, deletions, duplications, translocations, inversions, and fusions greater than 50 bp (Alkan, et al. 2011), represent a significant source of genetic variation within populations (Sudmant, et al. 2015). For example, in humans, two individuals may differ at 0.1% of their genomes when measured by SNVs, but by up to 1.2% due to SVs (Pang, et al. 2010). Furthermore, SVs are frequently associated with major phenotypic changes. The textbook example of the peppered moth (*Biston betularia*) which underwent a color change from white to black in response to coal pollution and bird predation during the industrial revolution was driven by a ∼22Kb SV that altered expression of the *cortex* gene (Van’t Hof, et al. 2016). Other SVs have been linked to disease phenotypes (Weischenfeldt, et al. 2013) for multiple human traits including cancers (Li, et al. 2020) and autism (Brandler, et al. 2018), while in pathogen populations SVs often play roles in drug resistance and immune evasion (Nair, et al. 2007; Nair, et al. 2008; Cheeseman, et al. 2016).

Despite their clinical and evolutionary significance, large SVs are often overlooked in population genetic studies of non-model organisms. This is primarily due to challenges in sequencing and genotyping. SVs are typically larger than the reads generated by short-read sequencing technologies, requiring indirect inference from secondary features of mapped short reads (Gong, et al. 2021). While these methods have achieved some success, they can suffer from relatively low accuracy and precision (Sibbesen, et al. 2018). Consequently, nearly 70% (n=107,590) of SVs maybe missed (Ebert, et al. 2021), representing a significant blind spot in the study of within-species variation.

Rapid improvements in long-read sequencing technologies by Oxford Nanopore Technologies and Pacific Biosciences now allow generation of reads over 100kb in length (Logsdon, et al. 2020). However, these methods are still expensive relative to short read technologies, and require large amounts of intact DNA from fresh tissue for best results. The central aim of this study was to evaluate the efficacy of pooled long read sequencing for rapid, cost-effective characterization of SVs segregating within populations. Use of pools has several advantages. First, single individuals may provide insufficient DNA for nanopore sequencing from fresh tissue; but see Kim, et al. (2023), Lee, et al. (2021), and Dadzie, et al. (2024) which show that high quality data may be generated from DNA generated by whole genome amplification. As an alternative, sufficient intact DNA can easily be obtained from pools of individuals. Second, use of pools allows cataloging of SVs in large numbers of individuals. Finally, analysis of pools provides information on allele frequency of SVs within populations. Pooled short read sequencing is now widely used for evaluating allele frequency of SNVs (Schlötterer, et al. 2014; Czech, et al. 2024), but has not been employed for long-read sequencing and SV characterization.

We focus on trematodes in the genus *Schistosoma* which includes blood-feeding flukes that parasitize humans and other mammalian hosts. These parasites are diploid, with a ZW sex determination and ∼400Mb genomes (Berriman, et al. 2009; Protasio, et al. 2012; Buddenborg, et al. 2021). *Schistosoma* larvae (miracidia) infect snail intermediate hosts, where they clonally reproduce to generate motile larvae (cercariae, second-stage larvae). These penetrate the skin of the mammalian host, migrate through the circulatory system, and settle in the mesenteric veins. There, mated female worms release hundreds of eggs per day (Moore and Sandground 1956). Eggs are expelled in feces or urine, restarting the life cycle (Nelwan 2019). However, many eggs also become lodged in various organs, resulting in pathology. Schistosomiasis affects over one hundred million people in the Global South, with the majority of the burden in sub-Saharan Africa (Steinmann, et al. 2006).

*S. mansoni* is particularly amenable to laboratory maintenance (Hackett 1993), and several laboratory populations exist that vary in their geographic origin or vary in biomedically important phenotypes. The genetics of wild and laboratory schistosomes have been extensively studied using SNVs (Berriman, et al. 2009; Protasio, et al. 2012; Clément, et al. 2013; Le Clec’h, et al. 2018; Chevalier, et al. 2019; Buddenborg, et al. 2021; Le Clec’h, Chevalier, Mattos, et al. 2021; Platt, et al. 2022; Vianney, et al. 2022) and previous work has shown that *S. mansoni* lab populations are diverse when measured at the single nucleotide level (Jutzeler, et al. 2024). However, little is known about genetic diversity at the level of SVs in lab or field populations.

Our study had three goals; (i) to catalogue SVs circulating in *S. mansoni* lab populations using long-read sequencing of pooled individuals as a proof-of-concept; (ii) to identify population specific SVs that may reflect adaptation (ii) to examine genome regions known to underlie parasite phenotypes for the presence of SVs. We have previously used genome-wide association (GWAS) and linkage studies to map quantitative trait loci associated with schistosome phenotypes such as drug resistance or larval stage production (Anderson, et al. 2018; Le Clec’h, Chevalier, McDew-White, et al. 2021). These mapping studies typically use short-read sequence data that may miss large SVs. SVs present in quantitative trait loci (QTL) regions could be an overlooked source of variation contributing to parasite phenotypes, as is the case in other species (Chakraborty, et al. 2018). To accomplish these goals, we sequenced pools of parasites from five *S. mansoni* laboratory populations using to describe 17,446 SVs large SVs circulating in these populations. Our results show that SVs can impact up to 6.5% of the genome, including 56 genes, with some populations harboring fixed population-specific variants (n=168) that may have evolutionary significance. Moreover, we found that large SVs overlap with known QTLs associated with parasite phenotypes. These observations underscore the importance of SVs in shaping genetic diversity within schistosome populations and demonstrate how pooled nanopore sequencing can provide an efficient rapid approach for assessment of segregating SVs.

## Results

### Sequencing Results

We generated long read data with Oxford Nanopore Technologies (ONT) sequencing platform for five lab populations of *S. mansoni*; SmBRE, SmEG, SmOR and two populations recently selected from a single population SmLE-PZQ-ER and SmLE-PZQ-ES (Table 1). Instead of sequencing a single individual from each population, we generated libraries from 92 -152 pooled male and/or female adult worms per population. Pooling individuals allowed us to isolate larger quantities of high molecular weight DNA, which is essential for long-read sequencing. This approach also offers the advantage of identifying the presence and frequency of variants across multiple individuals, providing a more accurate reflection of the structural variants (SVs) present within each lab population compared to sequencing one or a few individuals at a time. The total amount of whole genomic DNA extracted from these pools ranged from 2 to 5 µg.

**Table 1.**
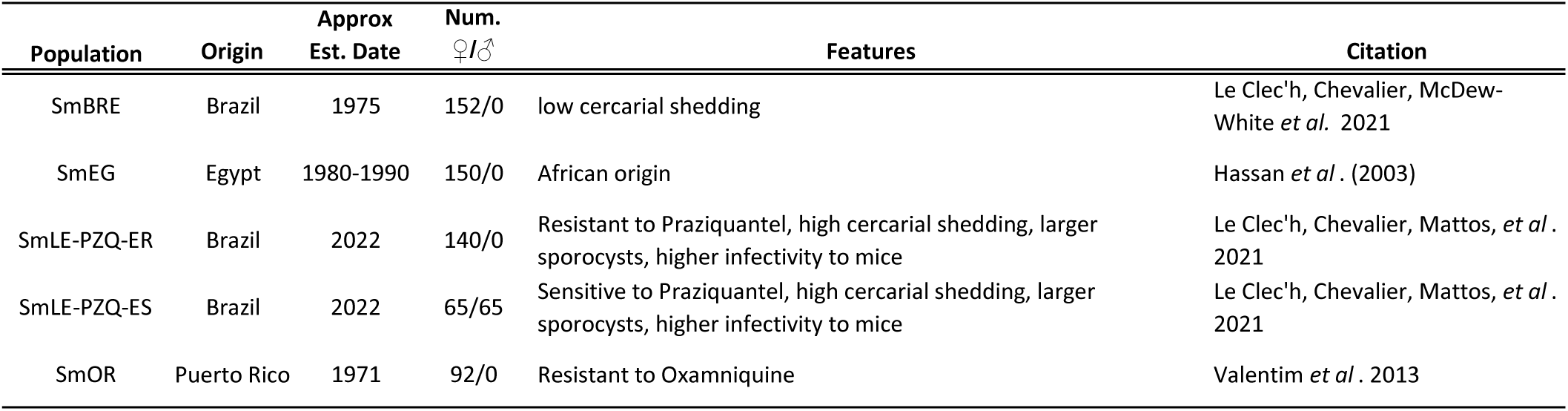
Populations Examined.

Across the five lab populations, we generated a combined total of 11.9 million reads, resulting in 66.7 gigabases of filtered data. The number of reads, genome coverage, and N50 values varied between populations, as shown in Table 2. N50 values ranged from 8,668 bp (SmOR) to 16,145 bp (SmBRE), with maximum mapped read lengths reaching up to 447 Kb (SmLE-PZQ-ES). Data quantity and quality was similar between the two methods, however, we prefer the Monarch kits, because they readily available. Mean genome coverage varied from 16.93× (SmLE-PZQ-ER) to 50.96× (SmLE-PZQ-ES). All sequence data is accessioned under NCBI BioProject PRJNA1206205.

**Table 2.**
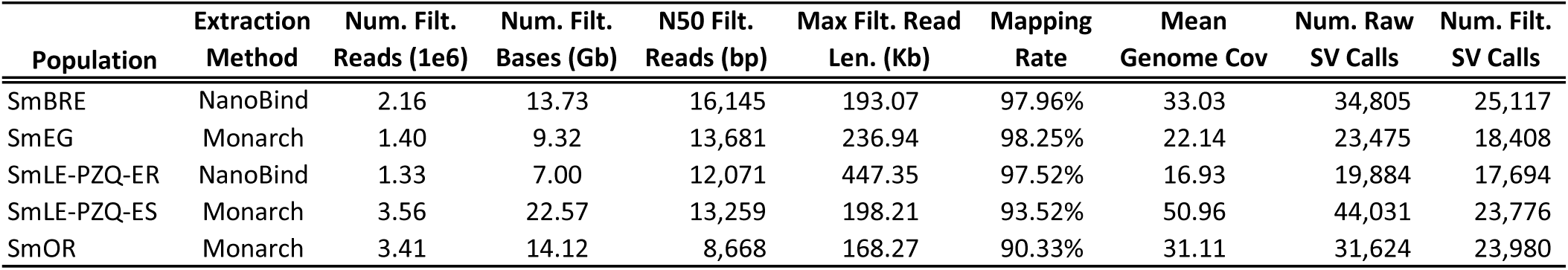
Summary of Long-read seqeunce data, genome coverage, and *de novo* SV identification.

| Population | Extraction Method | Num. Filt. Reads (1e6) | Num. Filt. Bases (Gb) | N50 Filt. Reads (bp) | Max Filt. Read Len. (Kb) | Mapping Rate | Mean Genome Cov | Num. Raw SV Calls | Num. Filt. SV Calls |
| --- | --- | --- | --- | --- | --- | --- | --- | --- | --- |
| SmBRE | NanoBind | 2.16 | 13.73 | 16,145 | 193.07 | 97.96% | 33.03 | 34,805 | 25,117 |
| SmEG | Monarch | 1.40 | 9.32 | 13,681 | 236.94 | 98.25% | 22.14 | 23,475 | 18,408 |
| SmLE-PZQ-ER | NanoBind | 1.33 | 7.00 | 12,071 | 447.35 | 97.52% | 16.93 | 19,884 | 17,694 |
| SmLE-PZQ-ES | Monarch | 3.56 | 22.57 | 13,259 | 198.21 | 93.52% | 50.96 | 44,031 | 23,776 |
| SmOR | Monarch | 3.41 | 14.12 | 8,668 | 168.27 | 90.33% | 31.11 | 31,624 | 23,980 |

### Identifying and Genotyping Structural Variants

We identified 153,819 raw structural variants (SVs) and indels across all populations during our initial round of genotyping, with counts ranging from 19,884 in SmLE-PZQ-ER to 44,036 in SmLE-PZQ-ES (Table 2). After applying a read depth filter (DP > 10), 4,821 SVs were removed, with the majority (43.5%, n = 2,095) being removed from SmLE-PZQ-ER, which had the lowest median genome coverage.

The combined impact of the DP > 10 and QUAL > 10 filters reduced the dataset from 153,819 to 108,975 total SVs (29.2% reduction). We applied these filters for two reasons. First, removing variants with low QUAL reduces false positive SV calls. Second, the next step in our analysis involved combining filtered variants across all populations. This means that a low-frequency variant removed due to a low QUAL score in one population might be at high frequency, have a high QUAL score, and be retained in another population. In such cases, the variant would still be included in downstream analyses.

The 108,975 variants from the five populations were merged into a single catalogue in a single step that combined overlapping variants and removed any remaining small indels (<50bp, n=70,216) leaving 17,565 large, unique SVs. A basic summary of the counts and sizes of each of the major SV types is shown in Table 3. We started with 17,565 variants, but dropped an additional 119 overlapping SVs that occurred at the same position in the genome, leaving 17,446 SVs. Of these 97.1% (16,935) were insertions and deletions, while 311 (1.8% were inversions and 69 (0.4%) were translocations (Figure 2). Combined, all SVs identified in these analyses impact 25.97 Mb representing 6.53% of the 397.9 Mb *S. mansoni* genome.

**Figure 1.**
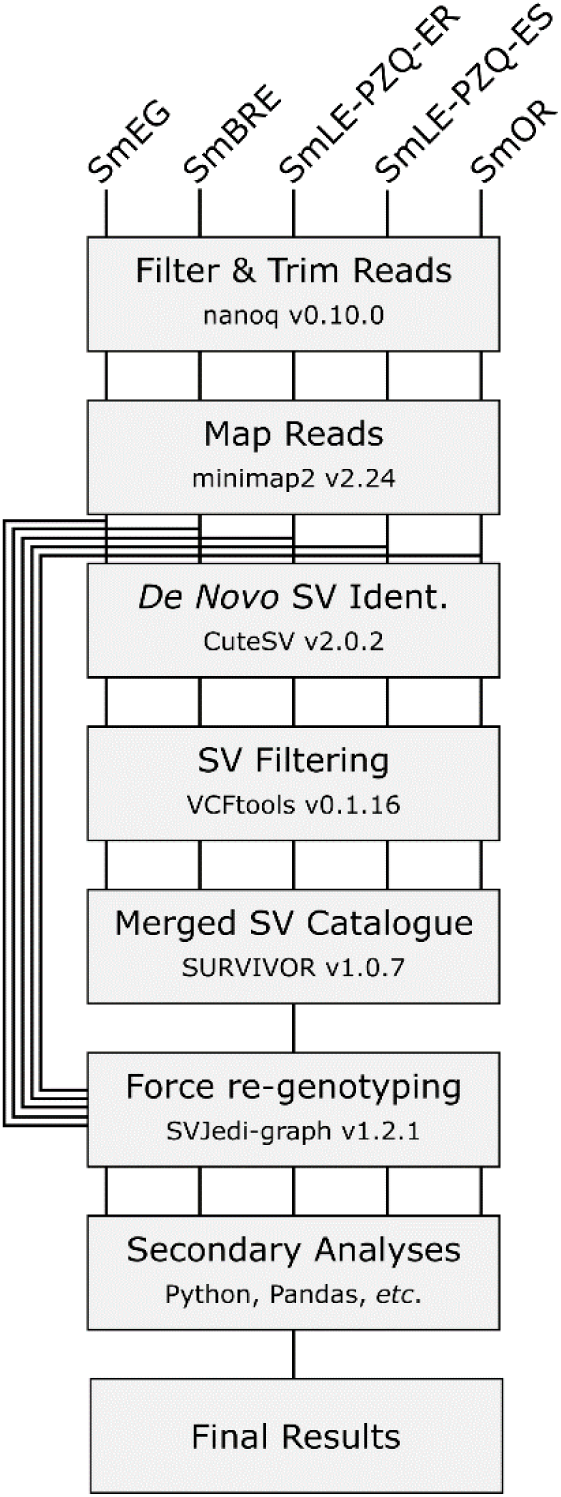
Bioinformatic Methods Overview. – Long read sequence data was generated for 5 *Schistosoma mansoni* populations in order to describe segregating structural variants (SVs). Initially we identified SVs in each population after filtering and mapping reads to the SM_V10 genome assembly. These initial SV calls were filtered and then merged into a single catalogue representative of all SVs within the S. mansoni populations we queried. We used that catalog of SVs to re-genotype the populations at each SV site in the catalogue.

**Figure 2.**
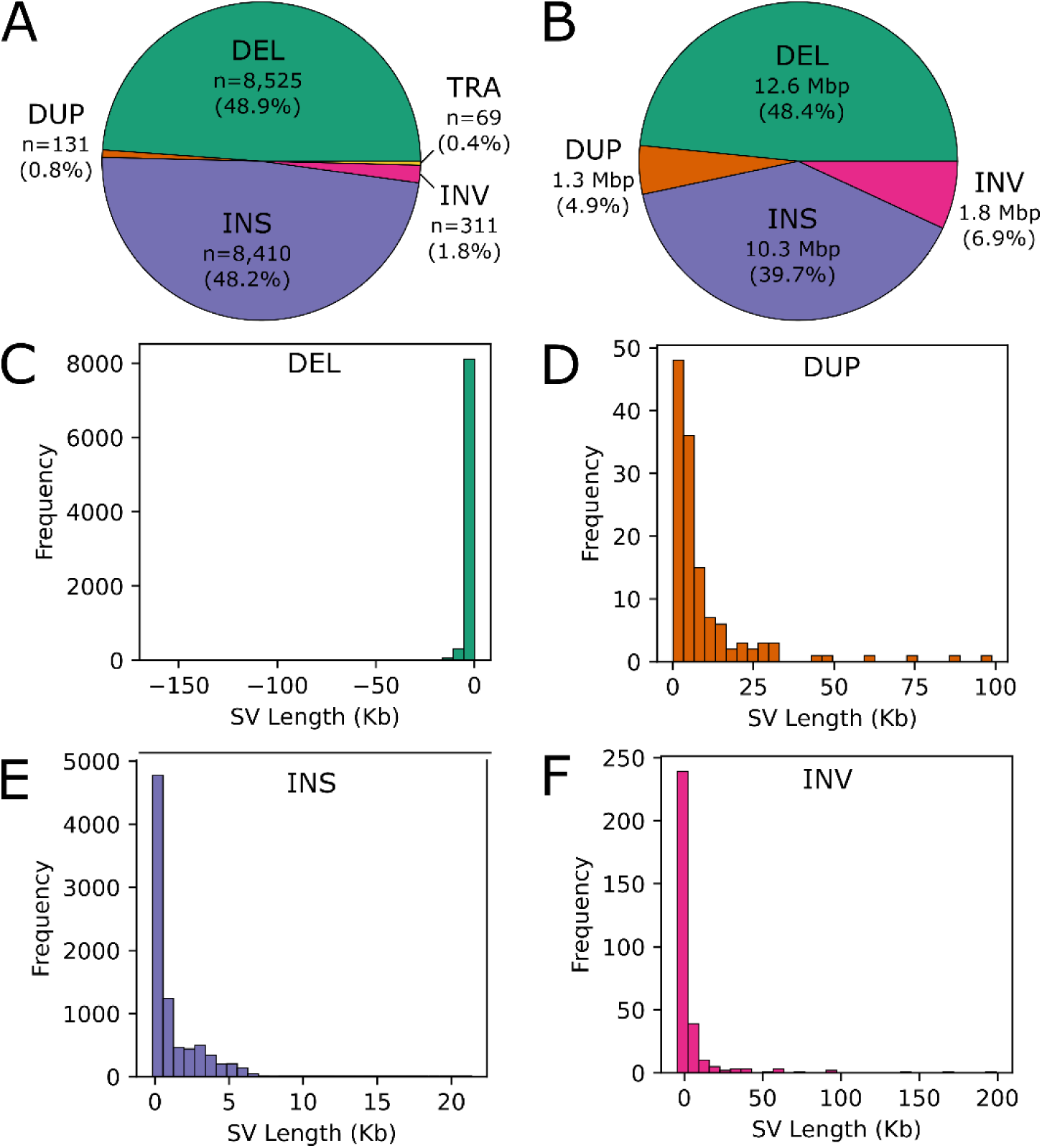
Abundance and size distributions of structural variants (SVs) in 5 *S. mansoni populations*. We examined five major types of SVs in total including; deletions (“DEL”), insertions (“INS”), inversions (“INV”), translocations (“TRA”) and duplications (“DUP”). (A) Most SVs were insertions or deletions, and only ∼3% were duplications, transversions, or translocations. (B) Similarly, the genomic impact in terms of bp affected by SVs is biased towards insertions and deletions. Breakpoints in translocations often occurred between chromosomes, or were on the same chromosome between telomeres so they were excluded from this subfigure. As a group, the size range for (C) deletions, (D) duplications, (E) insertions, and (F) inversions varied between 50bp-199,209 bp.

**Table 3.** Summary statistics for each of the major SV types after filtering and merging from all five lab populations.

|  | Count | Mean (bp) | Median (bp) | Min (bp) | Max (bp) | Total bp | Count |
| --- | --- | --- | --- | --- | --- | --- | --- |
| Deletions (DEL) | 8,525 | -1,475 | -441 | -161,688 | -63 | 12,576,321 | 8,525 |
| Duplications (DUP) | 131 | 9,798 | 4,504 | 73 | 98,879 | 1,283,568 | 131 |
| Insertions (INS) | 8,410 | 1,227 | 427 | -179 | 21,343 | 10,320,844 | 8,410 |
| Inversions (INV) | 311 | 5,745 | 864 | -4,424 | 199,209 | 1,786,834 | 311 |
| Translocations (TRA) | 69 | Na | Na | Na | Na | Na | 69 |

We re-genotyped each population using the catalogue of 17,446 SVs that span entire genome including autosomes, sex chromosomes, and the mitochondria. Allele frequencies were calculated for each SV at the population level (Figure 3A). The alternate allele frequency distributions were similar between populations and variant types. Fixed variants, where the alternate allele is at 0% or 100% frequency, were particularly abundant, representing 23.5% to 50.2% of any SV type and population combination. Segregating variants were evenly distributed at lower frequencies across each SV type and population combination.

**Figure 3.**
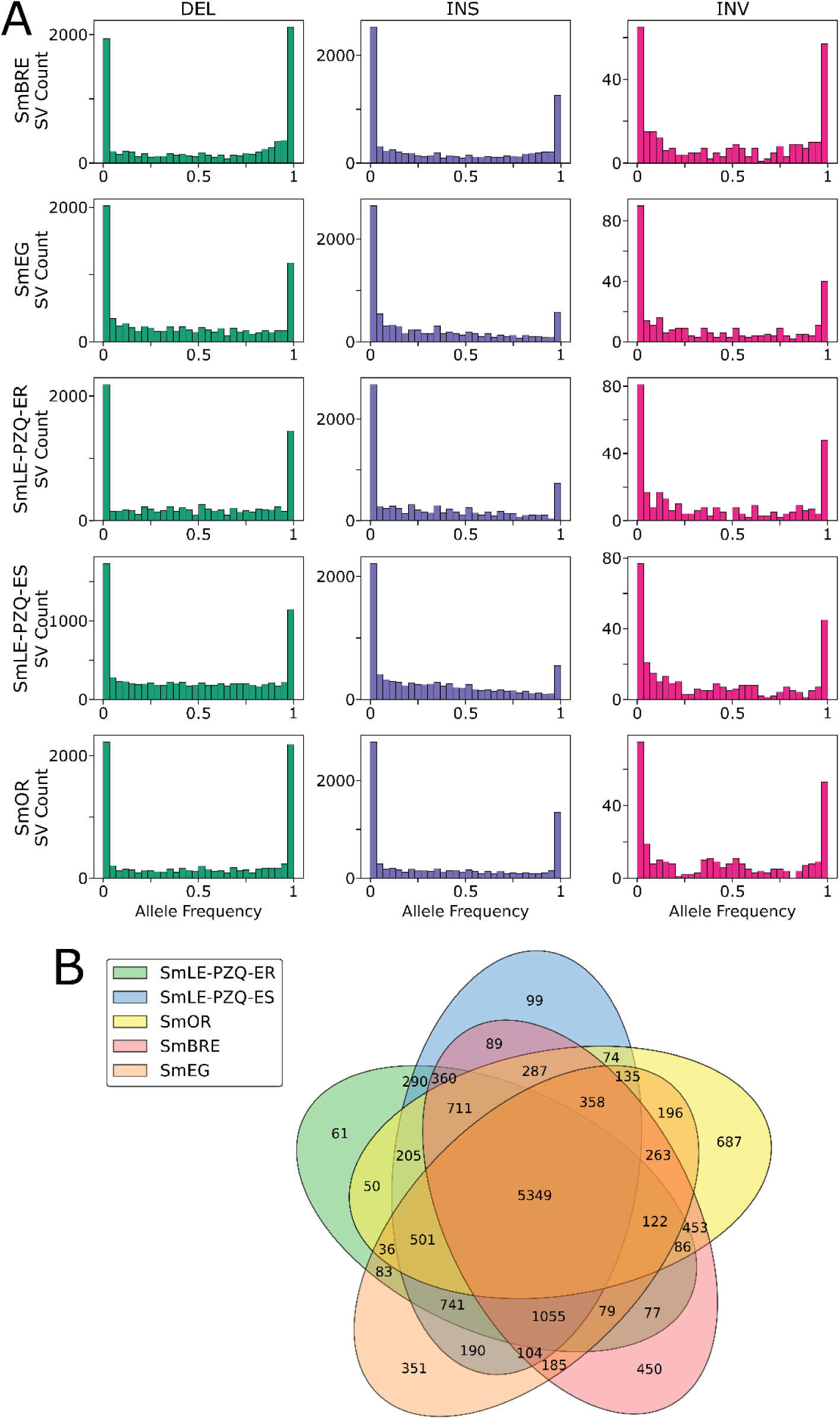
SV distributions in S. mansoni populations. (A) Allele frequencies (AF) distributions for each lab population of *S. mansoni* broken down by SV type. The AF distributions were relatively consistent for each population and SV type. Most SVs were at-or-near fixation (AF < 0.05 | AF >0.95). All SVs and allele frequencies are called relative to the reference. For example, a deletion specific to the reference genome is identified as a fixed insertion in the other populations. This leads to the bias of high frequency SVs shown above. (B) A Venn diagram showing the distribution of SVs among 5 lab populations of *S. mansoni*. SVs were scored as “present” if they were present in the population with an alternate allele frequency >5%.

### SV distributions among populations

We examined the distribution of SVs among each of the five lab populations. Determining absence of an SV is problematic because each population differed in read depth and genome coverage. This affects precision and our ability to confidently state that a variant is missing from a population. To circumvent this issue, we treated all variants at <5% in a population to be “low frequency” rather than absent. Using this criterion, we were able to isolate 1,648 SVs (11.3%) that were at >5% in a single population (Figure 3B). By comparison 5,349 (36.8%) of all autosomal SVs were at >5% frequency in all populations. While most population specific SVs were at low to moderate frequencies, we found 168 population specific SVs that are at-or-near fixation (>95% alternate allele frequency). All the population specific SVs were found in three populations; 114 in SmOR, 46 in SmBRE, and 8 in SmEG (Figure 4, Supplemental Table 1). We did not identify any SVs that were specific to the SmLE-PZQ-ER and SmLE-PZQ-ES, consistent with the recent divergence of these two populations (Le Clec’h, Chevalier, Mattos, et al. 2021).

**Figure 4.**
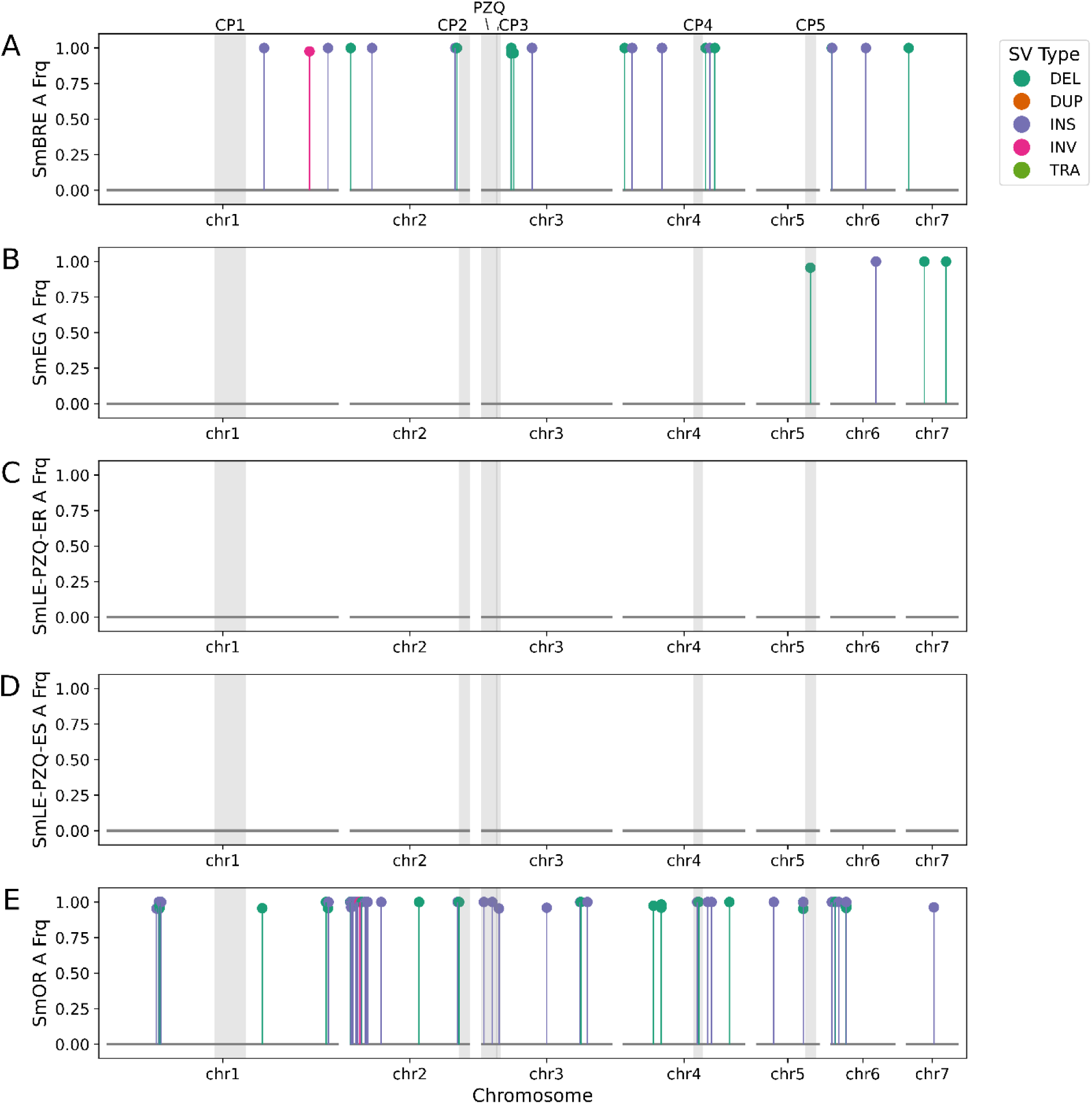
Population specific, structural variants (SV) distribution across the genome. Population specific variants are at >95% in a single population and <5% in the other four other populations. The distribution of population-specific SVs is not uniformly distributed across the genome. Grey boxes indicate parasite QTLs identified in previous studies and includes 5 QTLs associated with cercarial production “CP” and a praziquantel resistance QTL (PZQ). We did not identify any population specific SVs in SmLE-PZQ-ER and SmLE-PZQ-ES. These populations were only recently established from an SmLE source population.

We examined the relationships between parasite populations using SV allele frequencies using a principal component analysis (PCA). To focus on autosomal variants and avoid potential biases from skewed sex distributions, we removed 3810 variants located on the Z chromosome, WSR, and mitochondria, resulting in a final dataset of 13,656 SVs. Nearly two-thirds of the variation in the PCA (Figure 5) was explained by the first two principal components: principal component 1 (36.79%) and principal component 2 (27.88%). The populations were evenly distributed in PC space, with the exception of SmLE-PZQ-ER and SmLE-PZQ-ES, which nearly overlapped. This overlap is not unexpected, as these two populations were only recently derived from a common SmLE-PZQ-R progenitor stock (Le Clec’h, Chevalier, Mattos, et al. 2021).

**Figure 5.**
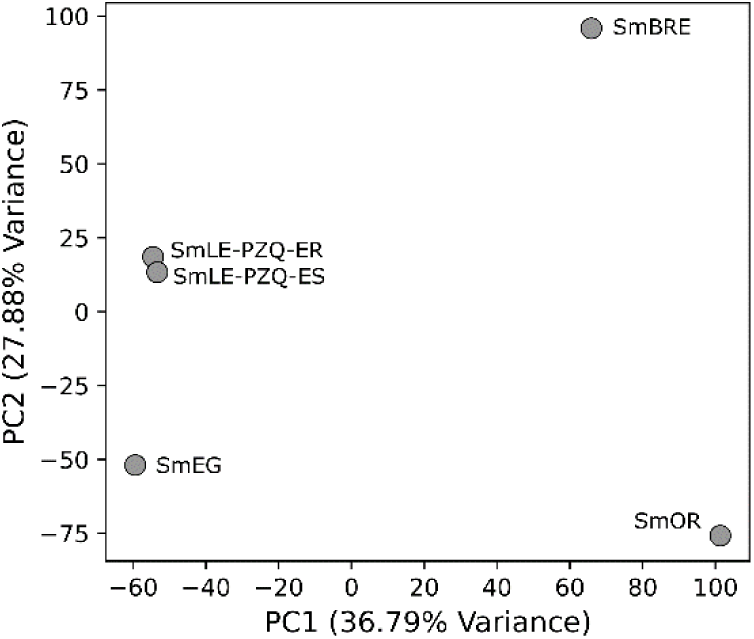
Population structure of five *S. mansoni* lab populations using allele frequencies from 17,446 SVs. SVs were scored as present or absent in a population if the allele frequency was >5%.

### SV impact on gene regions

We examined structural variants (SVs) in coding sequences and regulatory regions which have the potential to affect parasite phenotypes (Figure 6). We identified 168 population-specific SVs, of which 70 (41.7%) overlap with 56 different genes (Supplemental Table 1). Most of these variants are located in intronic regions (n=50); however, we found population-specific variants affecting six genes within either the UTR or coding sequence. These variants include one inversion, one deletion, and four insertions. The inversion spans approximately 30 Kb (SM_V10_2:3,848,692-3,878,697) and contains two genes (Smp_327700 and Smp_327710). Both are identified as secreted proteins by WormBase ParaSite (WBPS19), but little is known about their expression profiles or functions. Similarly, there is limited information available on Smp_158710 which contains a deletion that removes part of an intron and 41 bp of an exon3 (SM_V10_6:6,022,561-6,022,739) in SmOR. Smp_315070, (a SERPIN domain-containing protein) which contains a pair of 577 and 2,293 bp insertions in the 5’ UTR of SmOR. Smp_315070, is expressed at low levels in many different tissue types (Wendt, et al. 2020). Smp_342090, a low-density lipoprotein receptor class A, contains a 364 bp insertion in the 5’ UTR in SmBRE. This gene is highly expressed in most types of neuronal cells (Wendt, et al. 2020). The final gene, Smp_076960, encodes a C2 NT-type domain-containing protein that is expressed in most major cell types (Wendt, et al. 2020). There is a 419 bp insertion in the 3’UTR of this gene in SmOR.

**Figure 6.**
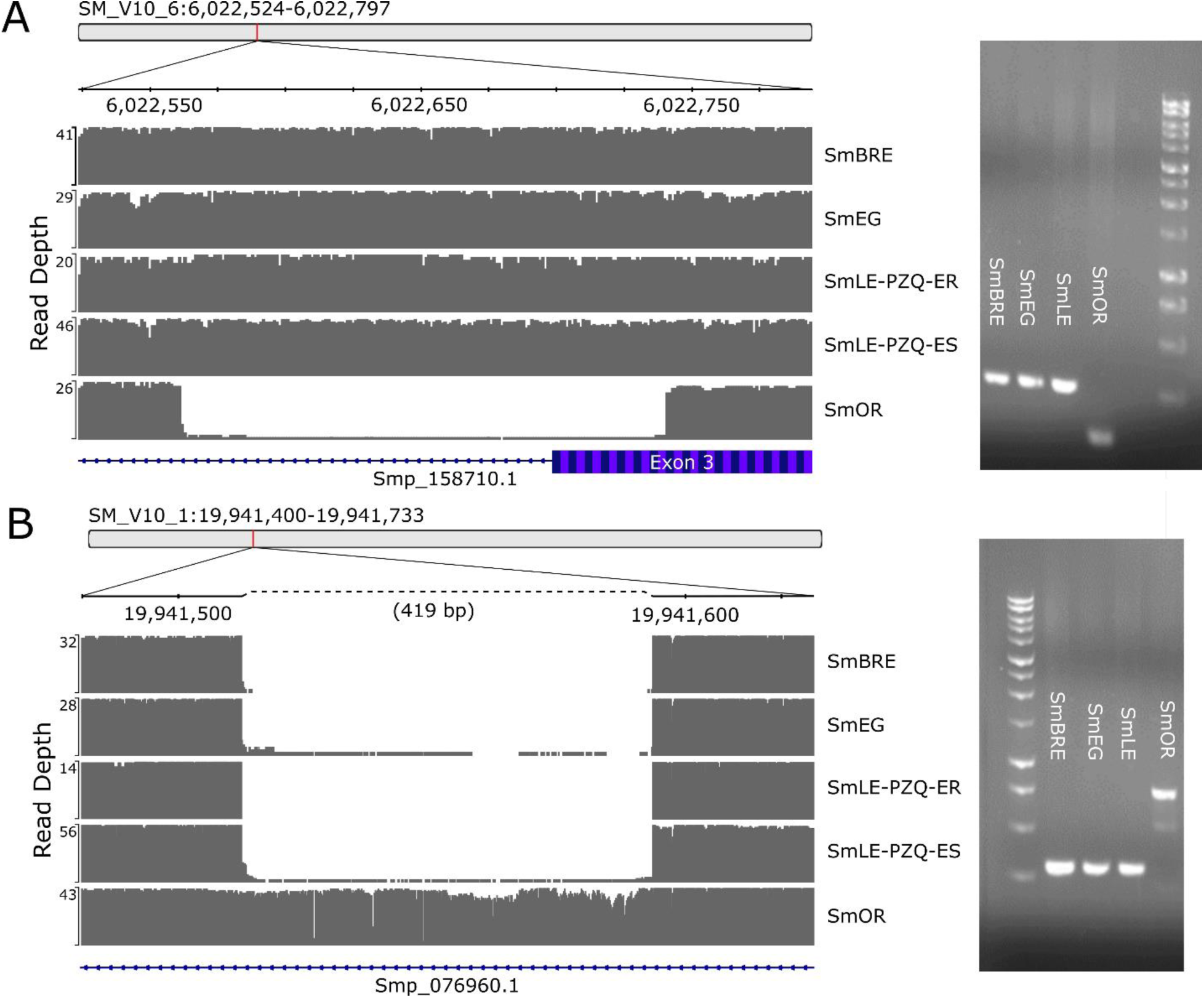
Examples of structural variants (SVs) within genic regions. (A) A 178 bp deletion in a gene (Smp_158710.1) with unknown function is specific to the SmOR population removes a portion of exon 3. (B) A 419 bp insertion within the 5’ UTR of a C2 NT-type domain-containing (Smp_076960.1). Both SVs were verified using PCR.

### SV within known parasite QTLs

Previous work has established six quantitative trait loci (QTLs) associated with either praziquantel drug resistance (Le Clec’h, Chevalier, Mattos, et al. 2021; Chevalier, et al. 2024) and cercarial production (Le Clec’h, Chevalier, McDew-White, et al. 2021). We converted the coordinates of these loci to the current SM_V10 assembly and examined the structural variants (SVs) within these regions, focusing on variants with strong allele frequency differences (>95%) between the populations analyzed in the original studies (Table 4).

**Table 4.**
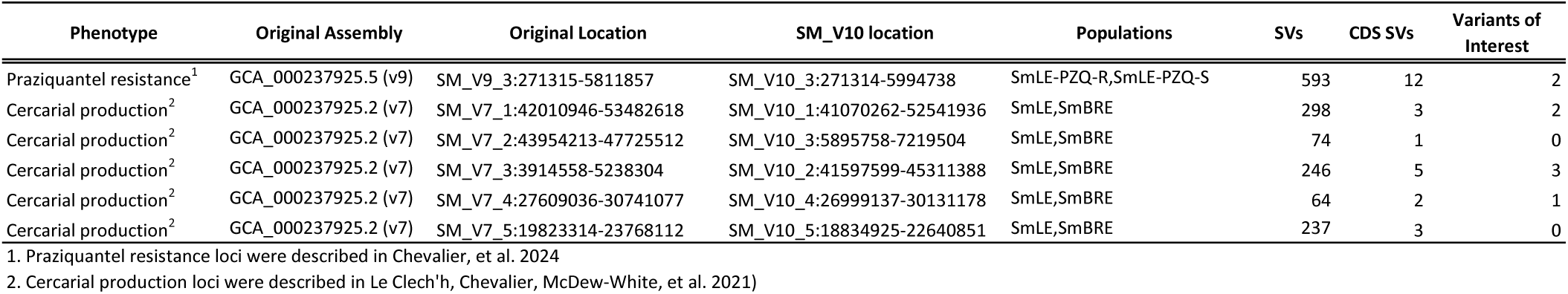
Quantitative trait loci (QTLs) associated with parasite phenotypes and their locations in the current *S. mansoni* genome assembly.

| Phenotype | Original Assembly | Original Location | SM_V10 location | Populations | SVs | CDS SVs | Variants of Interest |
| --- | --- | --- | --- | --- | --- | --- | --- |
| Praziquantel resistance<sup>1</sup> | GCA_000237925.5 (v9) | SM_V9_3:271315-5811857 | SM_V10_3:271314-5994738 | SmLE-PZQ-R,SmLE-PZQ-S | 593 | 12 | 2 |
| Cercarial production<sup>2</sup> | GCA_000237925.2 (v7) | SM_V7_1:42010946-53482618 | SM_V10_1:41070262-52541936 | SmLE,SmBRE | 298 | 3 | 2 |
| Cercarial production<sup>2</sup> | GCA_000237925.2 (v7) | SM_V7_2:43954213-47725512 | SM_V10_3:5895758-7219504 | SmLE,SmBRE | 74 | 1 | 0 |
| Cercarial production<sup>2</sup> | GCA_000237925.2 (v7) | SM_V7_3:3914558-5238304 | SM_V10_2:41597599-45311388 | SmLE,SmBRE | 246 | 5 | 3 |
| Cercarial production<sup>2</sup> | GCA_000237925.2 (v7) | SM_V7_4:27609036-30741077 | SM_V10_4:26999137-30131178 | SmLE,SmBRE | 64 | 2 | 1 |
| Cercarial production<sup>2</sup> | GCA_000237925.2 (v7) | SM_V7_5:19823314-23768112 | SM_V10_5:18834925-22640851 | SmLE,SmBRE | 237 | 3 | 0 |

1. Praziquantel resistance loci were described in Chevalier, et al. 2024 2. Cercarial production loci were described in Le Clech'h, Chevalier, McDew-White, et al. 2021)

In the single QTL associated with praziquantel drug resistance, we found two variants of interest. While these variants are strongly differentiated between SmLE-PZQ-ER and SmLE-PZQ-ES, they are also found circulating in other *S. mansoni* populations at frequencies ranging from 0 to 100%. Notably, two variants, DEL.SM_V10_3:2,969,428-2,973,340 and DEL.SM_V10_3:3,194,040-3,194,564, impact the intronic regions of Smp_345310 (Transcription factor SOX-13), and Smp_307630 (Zinc finger, TFIIS-type; Pol I subunit A12, C-terminal zinc ribbon), respectively. Among the five QTLs associated with cercarial production, two SVs exhibited a >95% frequency difference between SmBRE and the SmLE populations: a 6,146 bp insertion at SM_V10_5:21925347 between Smp_330830 and Smp_086690 (Protein kinase domain; Serine/threonine-protein kinase, active site; Protein kinase, ATP binding site) and a 76 bp insertion (INS.SM_V10_2:41902922). In these cases, we used the mean frequency of these SVs in SmLE-PZQ-ER and SmLE-PZQ-ES, which were derived from SmLE.

#### Repeats and SVs

*De novo* searches of the genome identified 985 families of repetitive sequences. We could assign only 228 of these to known transposable element families and 76.9% (n=757) were unknown repetitive sequences. More than half (53.3%) of the genome was derived from repetitive sequences, with the largest class of elements falling into the unknown category (27.05%) followed by retrotransposons (24.39%; Figure 7). Bov-B RTEs and L2s are the most common families of described retrotransposons in the genome. By comparison 84.01% of SV alleles are derived from repetitive sequences and the difference between the two categories (whole genome vs. SV alleles) is driven almost entirely by the expansion by the Gypsy family of long-terminal repeats (LTRs). These are present in 4.1% of the genome but in 23.86% of the SV alleles. The overall repeat content in the SV alleles is significantly greater than in the genome as a whole (two-proportion z-test: *p*=0.0, *Z-score*=2,916). The strong association between repeats and SVs provides a further limitation on our ability to score SVs with short-read Illumina data, which cannot be effectively mapped in repeat regions.

**Figure 7.**
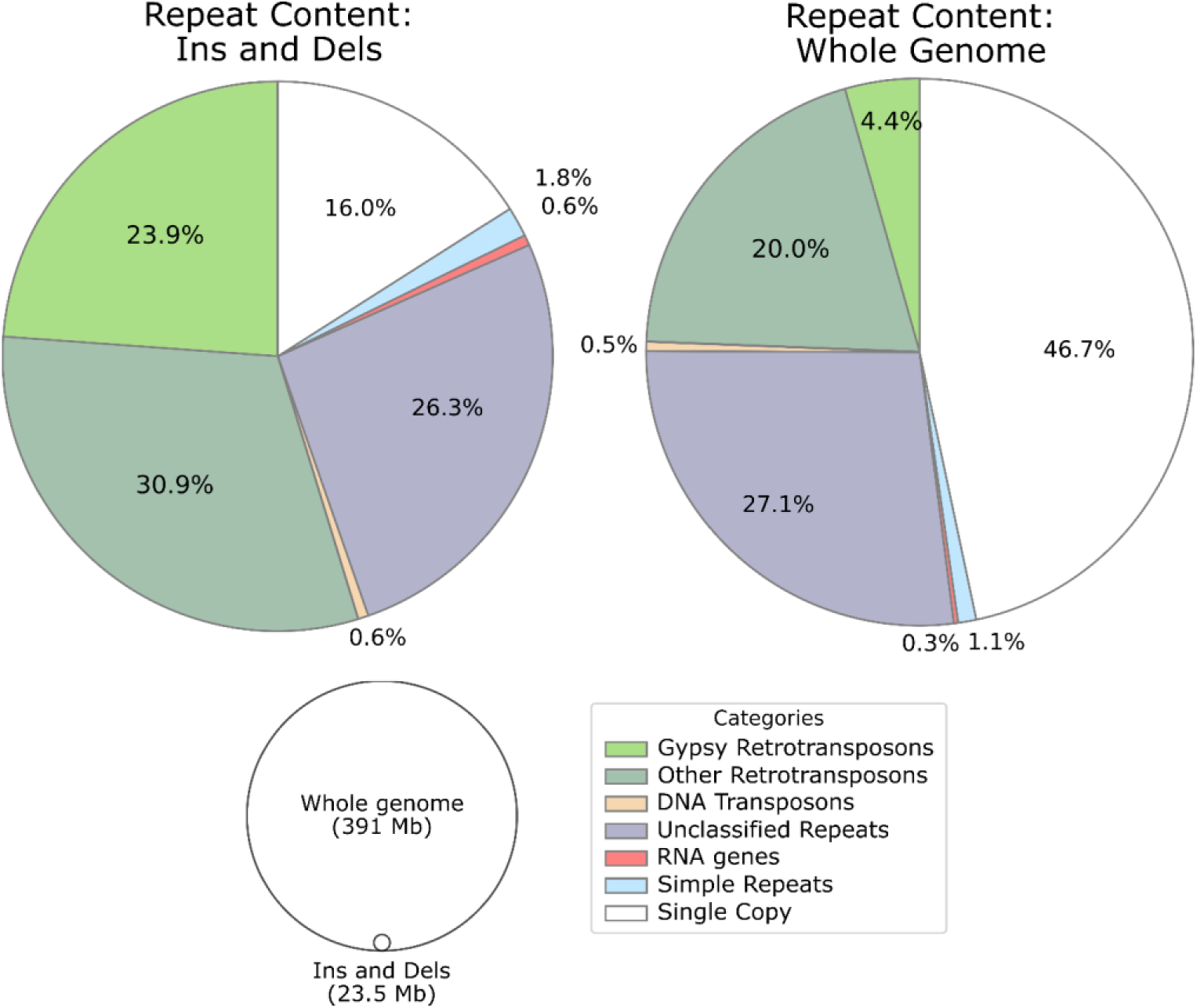
Structural variants (SVs) are frequently associated with repetitive sequences. Around half of the *S. mansoni* genome is derived from repetitive sequences, but 84% of SVs are either caused by, or found within repetitive regions of the genome. The difference, appears to be associated primarily with retrotransposon transposable elements (TEs), primarily those in the Gypsy family.

### SV validations

We used PCR methods to validate SVs from individuals within each population, focusing on population specific SVs found within genes or under QTL peaks. These included 11 insertions, 12 deletion and 2 inversions. We generated primers (Supplemental Table 2) to successfully amplify 19 of 25 loci. This included 12 population specific variants, 2 coding variants, and 5 variants in QTL regions. In general, the PCR results were consistent with the expected (Table 5; Supplemental Figure 3). We were able to validate SVs with 90% accuracy, 87% sensitivity and 93% sensitivity.

**Table 5.** PCR validation of Structural variants in SmBRE, SmEG, and SmOR.

|  | *n* | False<br>Neg | True<br>Neg | True Pos | False<br>Pos | Accuracy | Sensitivity | Specificity |
| --- | --- | --- | --- | --- | --- | --- | --- | --- |
| Pop-Specific | 12 | 2 | 44 | 20 | 0 | 97% | 91% | 100% |
| QTL | 5 | 2 | 6 | 10 | 2 | 80% | 83% | 75% |
| Genic | 2 | 0 | 8 | 4 | 0 | 100% | 100% | 100% |
| Overall | 19 | 6 | 63 | 34 | 5 | 90% | 85% | 93% |

Accuracy = (TP + TN)/(TP + TN + FP + FN), Sensitivity = TP/(TP + FN), Specificity = TN/(TN + FP)

## Discussion

The availability of long read sequencing is rapidly changing our ability to characterize SVs (Mahmoud, et al. 2019). In this study, we used long read sequencing to identify structural variants in five laboratory populations of schistosome worms with the goals of cataloguing large SVs in *S. mansoni* lab populations and identifying variants that may impact parasite phenotypes. By sequencing parasite pools, rather than individual parasites, we aimed to evaluate frequencies of SVs segregating within each population.

### Challenges in identifying low frequency variants in pooled samples

Pooled sequencing using short read, lllumina approaches is now widely used rapid, cost effective determination of SNP allele frequencies within populations (Lynch, et al. 2014). Comparable methods for measuring SV frequencies using nanopore long read sequences from populations could also be extremely valuable. However, identifying SVs from pools of individuals presents challenges compared to genotyping single individuals (Layer, et al. 2014). We initially identified 19,884 - 44,031 SVs for each of the five lab populations. This was reduced to 17,446 SVs after multiple rounds of filtering and merging into a single set of consensus variants. We outline several limitations and opportunities for analysis of such data.

### Limitations on calling low frequency SVs

Most SV tools, and quality metrics like QUAL and GQ, are designed to identify germline mutations that exist as homozygous or heterozygous variants. Pools of individuals defy these expectations since SVs can be present at frequencies ranging from 0-100% within a population compared to the 0%, 50%, and 100% expected in a single diploid individual. QUAL scores assess the confidence that a SV exists at a particular site. In our dataset, even moderate frequency (<27%) variants did not have QUAL scores higher than 10 indicating 90% confidence in a call (Supplemental Figure 1). This means that a low-to-moderate frequency variant in population “A” is only present in the final catalogue if it is also present at relatively high frequency in another population(s). As a result, our catalogue underreports low-to-moderate frequency variants in schistosomes populations.

### Sequence coverage and SV call accuracy

Genome coverage also contributes to accuracy in low frequency SV calls (Yang 2020). The number of SVs was positively correlated with genome coverage (Supplemental Figure 2) both before (*R*^2^=0.97) and after filtering (*R*^2^=0.77). Ideally, a population should be sequenced to a depth greater than the number of individuals (genomes) present in the pool to avoid missing low frequency variants. For example, in simulated pools of cancer cells, an SV at 20% allele frequency is only detected 31% of the time at 10x coverage (Layer, et al. 2014). We estimate that by sequencing 100 pooled worms from the SmLE-PZQ-ES population to 51x coverage, the lowest frequency variants we can score with 95% confidence is 5%. By contrast, we can only be confident in sampling variants present at >15% in SmLE-PZQ-ER due to lower total genome coverage (17x). Our inability to sample SVs at low coverages is compounded by the fact that low frequency variants are harder to distinguish from false positive and false negative calls using standard quality metrics.

### Long read sequencing of pooled individuals can be used to identify large SVs

By pooling parasites from each population, we were able to quickly survey SVs from five *S. mansoni* populations. Even with the limitations discussed above, we were able to identify 17,446 SVs that were segregating within the lab populations. Combined, these SVs, impact ∼26 Mb representing 6.5% of the genome. This number is almost certainly an underestimate since low frequency variants were filtered from the final SV catalogue.

SVs can be used as markers to understand relationships between populations. The majority of SVs where shared between ≥4 populations (n=8,096; 59%) compared to only 1,648 (12%) lineage-specific SVs. We used SV allele frequencies to project the populations in genotypic space using a PCA (Figure 7). While we are limited to 5 data points, the patterns are consistent with the history of each lab population. SmLE-PZQ-ER and SmLE-PZQ-ES are derivatives of the SmLE population based on their susceptibility to the drug praziquantel (Le Clec’h, Chevalier, Mattos, et al. 2021). In the PCA, the two populations are effectively overlapping. Further, these two populations combined contain 160 unique variants while each of the other populations contain at least 350 unique, moderate-frequency (>5%) structural variants (Figure 3B).

### Population-specific structural variants can affect coding sequences

While the majority of SVs were shared between multiple populations, we found strong evidence of lineage-specific variants. These SVs are at-or-near fixation (>95%) in one population and less than 5% in the others. We identified 7,315 (41.9%) SVs that occur within, or span, genic regions. Of these we found 168 population-specific variants, but only five of these directly impact coding regions (CDS) of 6 genes.

Little is known about these 5 genes in terms of gene function or expression. Smp_327700 and Smp_327710 are secreted proteins found in a 30Kb inversion, Smp_315070 (a SERPIN domain-containing protein), Smp_158710, and Smp_342090 contain insertions or deletions. Each of these SVs is specific to the SmOR population with the exception of Smp_342090 which is restricted to SmBRE. The final gene, Smp_076960 contains a 419 bp insertion at SM_V10_1:19,941,568 in the 5’ UTR of the C2 NT-type domain-containing protein. This variant is specific to the SmOR population. This gene appears to be ubiquitously expressed throughout the parasite’s life cycle, in both sexes, and is not localized to any specific region (Harris, et al. 2020). Gene expression of Smp_076960 is significantly impacted by the histone-modifying enzyme GSK343 (Pereira, et al. 2018). Additionally, an ortholog of this gene is associated with abnormal sleep recovery and lethargy in *C. elegans* (WBPhenotype:0001524; Huang, et al. 2017). While it is difficult to directly associate any of these variants directly with a phenotype, these results show that SVs likely to disrupt transcription are found within genes and show very different distribution in the populations examined.

Population-specific variants are not randomly distributed throughout the genome (Figure 4; chi-square test p-value = 1.2e-180). Instead, we found 7 windows accounting for 28Mb of the genome (∼7%) that contains 109 of the 168 (64.9%) of all population-specific variants. Within these regions, population-specific variants occur every 3.3 Kb (mean) and outside of these regions SVs are distributed every 441.8 Kb (mean).

### High frequency structural variants are found in QTLs associated with parasite phenotypes

SVs have been associated with phenotypes in many species including horses (Bellone, et al. 2023), pigs (Zong, et al. 2023), goats (Guo, et al. 2021), humans (Hollox, et al. 2022), and others (Zhang, et al. 2016; Jeffares, et al. 2017; Guo, et al. 2020; Chen, et al. 2023). We have used both linkage mapping and association studies to identify the genetic basis of parasite phenotypes in *S. mansoni* (Anderson, et al. 2018; Le Clec’h, Chevalier, McDew-White, et al. 2021; Chevalier, et al. 2024). We investigated whether SVs were located within QTL regions for praziquantel resistance or release of larval parasites (cercarial shedding) from the intermediate host snails. We found population specific SVs of interest in four of six QTLs examined. The presence of these SVs does not necessarily imply they are the causative variants for the observed phenotypes, but these are potential candidates. A transient receptor potential channel gene (TRPM-PZQ) is thought to underlie the praziquantel resistance QTL on chr 3 (Le Clec’h, Chevalier, Mattos, et al. 2021; Park, et al. 2021). However, specific causative SNPs within this gene have not been located. It is possible that SVs outside of the TRPM-PZQ result in altered expression of this gene and modify PZQ-resistance. Interestingly there is a 524bp deletion in the intronic regions of Smp_345310 (Transcription factor SOX-13): this gene shows the strongest association with praziquantel resistance within the QTL region (Le Clec’h, Chevalier, Mattos, et al. 2021; Chevalier, et al. 2024).

Chevalier *et al*. (2024) used short read SV calling method to identify two large deletions tightly linked with the praziquantel QTL. These deletions were at SM_V10_3:2,775,001-2,900,000 and SM_V10_3:3,175,001-3,300,000; referred to here as the “upstream” and “downstream” deletions respectively. We did not recover either of these deletions in the nanopore analysis and only found one deletion that differentiates the two SmLE populations within these QTL regions (SM_V10_3:3,194,040-3,194,564). Mean read depth across the deletions is 0.07x (upstream) and 1.11x (downstream) and both are in the 0.03 and 0.06 percentile when compared to the rest of the genome sampled in 125kb windows. This is consistent with large deletions. By comparison, these same regions are in the 96.2 and 17^th^ percentiles in SmLE-PZQ-ES. Visual inspection of the read alignments is not consistent with an immediate break typically associated with deletions. Instead, genome coverage gradually changes over hundreds of base pairs in regions with high repeat. In the end it is likely that this combined with low coverage in SmLE-PZQ-ER in general and variation in pooled populations likely negatively impact SV calling in these regions.

### Repetitive sequences are strongly linked with SVs in S. mansoni populations

We estimate that more than half (53.3%) of the *S. mansoni* genome is derived from repetitive sequences. This is higher than estimates from previous work on *S. mansoni* (45%; Berriman, et al. 2009) and S. *haematobium* (43%; Young, et al. 2012). The difference between our analysis and Berriman et al. (2009) could be driven by many factors. For example, Berriman *et al*. (2009) used RepeatScout (Price, et al. 2005) to identify repetitive sequences. Here we used RepeatModer2 (Flynn, et al. 2020) which uses a combination of RepeatScout (Price, et al. 2005), RECON (Bao and Eddy 2002), LTRharvest (Ellinghaus, et al. 2008), and LTR_retreiver (Ou and Jiang 2018). Most of the repeats (27.1%) from our *de novo* analysis were unique and did not share significant homology or structural features for classification. Hence, more than 100Mb of the 400Mb *S. mansoni* genome is derived from repeats of unknown origin. Future work should look to provide a complete annotation (Platt, et al. 2016; Goubert, et al. 2022) and deposit the repeats in a public repository (ex. RepBase; Bao, et al. 2015) as a first step in understanding the impact of repetitive sequences on the evolutionary history of *S. mansoni*.

It is apparent that *S. mansoni* SVs are significantly enriched in repetitive sequences (two-proportion z-test (*p*=0.0, *Z-score*=2,916.7). While 53.3% of the genome is repetitive, 84% of SVs are caused by, or occur in repetitive sequences. The overrepresentation of repeat sequences in SVs is consistent with previous work (Domínguez, et al. 2020; Reis, et al. 2023; Quah, et al. 2024; Smeds, et al. 2024). Further, we note that highly, repetitive sequences larger than 150bp are difficult to unambiguously map to with short read sequence data. This implies that genotyping SVs with short read sequence could potentially miss many of the variants associated with repetitive sequences.

The increase in repeat-related SVs is driven almost entirely by Gypsy endogenous retrovirus which are present in 4.4% of the genome but 23.9% of SV alleles. Structurally, endogenous retroviruses contain a combination of gag, pol, and env genes flanked by two, long terminal repeats (LTRs). The LTRs may range between 100-1,000bp in length and are exact duplicates of each other. Because of their size and sequence similarity, LTRs are hotspots for non-homologous recombination which removes the internal regions of the ERV and one of the LTRs, leaving a behind a single LTR (Hughes and Coffin 2004). The disproportionate number of Gypsy LTRs associated with SVs suggests that they are actively mobilizing in the genome, creating insertions, and being partially removed via non-homologous recombination, generating the deletions.

### Conclusions

We used long-read sequencing of pooled parasite populations to identify and describe SVs within five *S. mansoni* lab populations. While these methods are limited in their ability to de novo detect low frequency variants from pooled samples, we were able to identify 17,446 SVs impacting 6.5% of the genome. Furthermore, we identified population-specific variants present at high frequencies and SVs that fall within known parasite QTLS for important phenotypic traits. Such population specific SVs may reflect the action of selection or determine phenotypic differences between population so are of particular interest. As a proof of concept, this study demonstrates the value of using long read sequencing and pooled samples to survey SVs within parasite populations. To ensure that the variation generated by SVs is properly accounted for, future work should aim to improve low frequency variant identification, to evaluate SV frequencies with field populations, and examine the impact of SVs on parasite phenotypes.

## Materials and Methods

### Parasite populations

We examined 5 laboratory populations of *Schistosoma mansoni* (SmBRE, SmEG, SmLE-PZQ-ER, SmLE-PZQ-ES, SmOR). These parasite populations differ in resistance to anthelmintic drugs, specificity to the snail intermediate host, and were initiated with founder individuals from different geographic regions (Table 1). These parasite populations are maintained by laboratory passage using hamsters as the vertebrate host (where sexual reproduction occurs) and *Biomphalaria spp*. snails as the intermediate host. Two of the populations, SmLE-PZQ-ER and SmLE-PZQ-ES, were derived from the same SmLE stock population less than 10 years ago (Le Clec’h, Chevalier, Mattos, et al. 2021). Fresh schistosome worms were obtained from hamster perfusions in the laboratory.

### DNA extraction and sequencing

We used two different DNA extraction methods for extracting HMW DNA and Nanopore sequencing of schistosome (Table 2). For the first method, we used the Monarch HMW DNA Extraction Kit for Tissue (NEB #T3060) and manufacturer recommended protocols to isolate DNA from 92-152 worms for the SmEG, SmLE-PZQ-ES, and SmOR populations. For the second extraction method we isolated DNA from SmBRE and SmLE-PZQ-ES populations using a developmental version of the Nanobind HMW DNA extraction kit being designed by Circulomics specifically for *C. elegans*. We transferred 100 mg of worms to 2 mL pre-chilled Protein LoBind microcentrifuge tube. The tube was centrifuged at 16,000 x g for 15 s. We discarded the resulting supernatant, added 20 μL Proteinase K and 150μl Buffer NL, and resuspended the worm pellet by vortexing 1-2 seconds prior to incubating on a ThermoMixer at 900 rpm and 55 °C for 1.5 h. After spinning the tube on a mini centrifuge for 1 s to remove liquid from the cap, we added 20 μL RNase A and incubated on a ThermoMixer at 900 rpm and 55 °C for 1 h. After spinning the tube on a mini centrifuge for 1 s to remove liquid from the cap, we added 40 μL Buffer ESB, pulse vortexed 5X, and incubated on ice for 5 min. We centrifuged the tube at 2,000g at RT for 10 min, and transferred 200 μL of the supernatant to a new 1.5 mL Protein LoBind microcentrifuge tube using a wide bore pipette, and discarded the tube containing the precipitate. We added a Nanobind disk to the lysate, followed by 250 μL isopropanol, and mixed 10 times by inversion, to precipitate DNA and allow binding to the Nanobind disk. The DNA is initially opaque on addition of isopropanol but becomes clear during subsequent washing steps. We mixed the tube on a tube rotator at RT for 15 min. We used the Magnetic rack handling procedure (PacBio PN 102-587-900) for washing the disc with buffers CW1, CW2. We eluted DNA with the appropriate volume of EB buffer for 7 days at 4°C. We used Qubit dsDNA BR Assay and Nanodrop to record DNA concentration and purity. Nanodrop readings should have A260/A280 in the range of 1.85–2.02 and A260/A230 in the range of 2.03–2.35. We used the Short Fragment Eliminator kit from PacBio (PN 102-582-400) to eliminate DNA below 25kb. We stored DNA at 4°C.

### Long read sequencing

A single microgram of high molecular weight genomic DNA was used for preparing libraries to be sequenced on the MinION. We prepared libraries with either the Genomic DNA by Ligation (SQK-LSK109) kit, or Ligation Sequencing V14 (SQK-LSK114) kit, loaded on flowcells with R9.4.1 or R10.4.1 chemistry. The sequencing runs lasted from 48-72 hours. Halfway through the run, we paused sequencing, washed the flow cells with EXP-WSH004-XL kit to unclog the pores, loaded more of the same libraries, and resumed sequencing.

### Genotyping Structural Variants

Bioinformatic analyses are summarized in Figure 1. We assessed long read quantity and N50s with n50 v1.4.2 (Telatin, et al. 2021). All reads were quality filtered and trimmed with nanoq v0.10.0 (Steinig and Coin 2022). We trimmed bases with a minimum Phred score of 7 (--min-qual 7), removed the first and last 25 bases (--trim-start 25, -- trim-end 25) and any reads that were less than 500 bp after trimming (--min-len 500). We then mapped the quality-filtered reads to the *S. mansoni* V10 genome assembly (SM_V10) with minimap2 v2.24 (Li 2016) using the “-x map-ont” parameter set. The SM_V10 assembly was accessed via WormBase ParaSite (accession: WBPS19; Harris, et al. 2020). Genome coverage was calculated with mosdepth v0.3.2 (Pedersen and Quinlan 2018). Reads were sorted with SAMtools v1.19 (Li, et al. 2009) for downstream analyses. SV detection and genotyping occurred in three steps; (1) an initial *de novo* SV query in each population, followed by (2) filtering and merging the population SV calls into a single catalogue, then each sample was (3) re-genotyped using the catalogue of merged variants.

### Initial SV Identification

SVs were identified for each individual population with an initial query using cuteSV v2.0.2 (Jiang, et al. 2020; Jiang, et al. 2022). We used the following parameters for SV identification: “--max_cluster_bias_INS 100 --diff_ratio_merging_INS 0.3 -- max_cluster_bias_DEL 100 --diff_ratio_merging_DEL 0.3 --min_mapq 20 --min_read_len 500 - md 100 -mi 100 --min_support 5 --max_size 200000 –genotype”. The combination of these parameters was chosen to identify larger SVs between 10bp-200Kb supported by at least 5 reads with a minimum mapping quality (MAPQ) of 20. These parameters are recommended for long read-based SV detection in the cuteSV documentation (https://github.com/tjiangHIT/cuteSV v0.19).

### SV Filtering and Merging

After the initial round of SV identification with cuteSV we explored various combinations of QUAL, GQ, and DP filters (discussed in the results) before choosing to remove variants with QUAL<10 and read depth (DP) <10 using VCFtools v0.1.16 (Danecek, et al. 2011). The filtered SVs from each population were merged into a single catalogue using SURVIVOR v1.0.7 (Jeffares, et al. 2017) and the following parameters: max_distance_between_breakpoints = 100, min_calls=1, sv_type=1, sv_strand=0, dist_estimate=0, and min_size=50. These parameters assured that variants must be present in at least one population and if variants are present in multiple populations they are only merged if they SV boundaries are within 100bp of one another, the SV is of the same type, and greater than 50bp.

### Final SV Genotyping and Secondary Analyses

The merged catalogue of SVs contains the higher quality SV calls present in one or more *S. mansoni* populations. We provided these variants to SVJedi-graph v1.2.1 (Romain and Lemaitre 2023) for forced variant calling at SVs with a minimum of 2 supporting reads (‘—minsupport 2’). We then converted the population VCF files to tables using the BCFtools v1.14 (Danecek, et al. 2011) query. The alternate allele frequency of each variant was calculated by dividing the alternate allele depth (AD) by total depth. Subsequent processing was managed with Pandas v2.0.3.

We used a principal component analysis (PCA) and Venn diagrams to examine how SV are distributed between the populations. To minimize artifacts caused by biases in the sex-distributions between populations, we removed any variants on the Z chromosome, WSR, and mitochondria to focus solely on autosomal SVs. A PCA was calculated for the first two principal components (PCs) using the PCA() function implemented in scikit-learn v1.2.0. To identify population specific SVs we used the venn() v0.1.3 package to generate Venn diagrams of the autosomal SVs. Rather than strictly examining presence or absence, we partitioned and categorized each SV as low frequency (<5%) or moderate-high frequency (≥5%). These cutoffs were chosen to minimize false negatives in low coverage populations where a SV may be present at low frequencies but wasn’t sampled in our sequence data. We used the Venn diagram to identify specific variants present at ≥5% in only one population. From this pool, we identified any variants with an allele frequency of ≥95% for downstream analysis. We refer to these SVs as lineage-specific as they are at-or-near fixation (≥95%) in one population but present in <5% in the others populations.

We identified Repeat families de novo using RepeatModeler v2.0.3 (Flynn, et al. 2020) to query the entire SM_V10 genome assembly. We used RepeatMasker v4.1.7 (Smit, et al. 2015) to quantify repeats in the whole genome, and in the SV alleles separately, using the de novo repeat families as a custom library (“-lib”) and the sensitive (“-s”) options. We compared the overall repeat content of each category, whole genome vs. SV alleles, using a two-proportion z-test. Our null hypothesis was that the overall repeat content of the SV alleles was the same as the rest of the genome.

We used BEDtools v2.31.1 (Quinlan and Hall 2010) to intersect the SV and gene annotations for the SM_V10 genome assembly. Previous work has established 6 QTLs associated with either praziquantel drug resistance (Le Clec’h, Chevalier, Mattos, et al. 2021; Chevalier, et al. 2024) or cercarial production (Le Clec’h, Chevalier, McDew-White, et al. 2021). We converted these coordinates to the current version of the *S. mansoni* genome by aligning the SM_V7 (GCA_000237925.2) and the SM_V10 assemblies with minimap2 v2.24 (Li 2016) and the “asm5” alignment preset parameter. Coordinates were converted between the assemblies using paftools.js liftover 2.28-r1209 (Li 2016).

### Validating Structural Variant Calls

We validated a subset of SVs by amplifying the target locus from individual worms in each population and scoring the presence or absence of the variants. For this dataset, we examined 26 population-specific SVs that were between 250-750bp. Primers for each locus were designed using BatchPrimer3 v2.6.1 (You, et al. 2008) with an optimum primer size of 20nt, melting temp of 59°C, and size range of 400-600 nt. We extracted DNA from individual worms from each population using Qiagen’s DNeasy Blood and Tissue kit and performed Whole Genome Amplification using Genomiphi V2 DNA Amplification kit from Cytiva. We amplified PCR products using a 25-cycle PCR reaction using TaKaRa Taq polymerase kit and ran 1µL of the PCR product on a 1% agarose gel (94 volts/for 60 minutes). Gels were visualized on the BIO-RAD ChemiDoc Touch Imaging System and scored as homozygous presence, heterozygous, or homozygous absence for each of the individual worms. We then compared these results to the population allele frequency to determine whether a variant was a true positive, false positive, true negative, or false negative to calculate accuracy, sensitivity, and specificity.

## Data Availability

We conducted analyses on compute nodes with a maximum of 1 terabyte of memory and 96 cores at the Texas Biomedical Research Institute’s high-performance computing cluster. All software and computing environments were managed with Conda v22.9.0. Initial analyses from quality filtering raw sequence data through the final SV genotyping steps were accomplished with a SnakeMake v7.25 (Mölder, et al.) workflow. Secondary analyses were conducted in a series of Jupyter v4.2.0 notebooks. All code, including environmental recipe files, notebooks, workflows, and shell scripts are available through GitHub (https://github.com/nealplatt/sch_man_ont) and are permanently accessioned with Zenodo (https://doi.org/10.5281/zenodo.14510995). All sequence data is accessioned under NCBI BioProject PRJNA1206205.

## Acknowledgments

We thank Sandy Smith and John Heaner for providing computational support through Texas Biomedical Research Institute’s High-Performance Computing Cluster. Frederic Chevalier, Winka Le Clec’h, Elisha Enabulele, and Kathrin Bailey provided parasite material and constructive feedback during the course of this project. This research was funded by the National Institute of Allergy and Infectious Diseases (NIAD R01 AI166049-01) and by a Texas Biomedical Research Institute Forum Grant 20-04866.

---

**Supplemental Figure 1.**
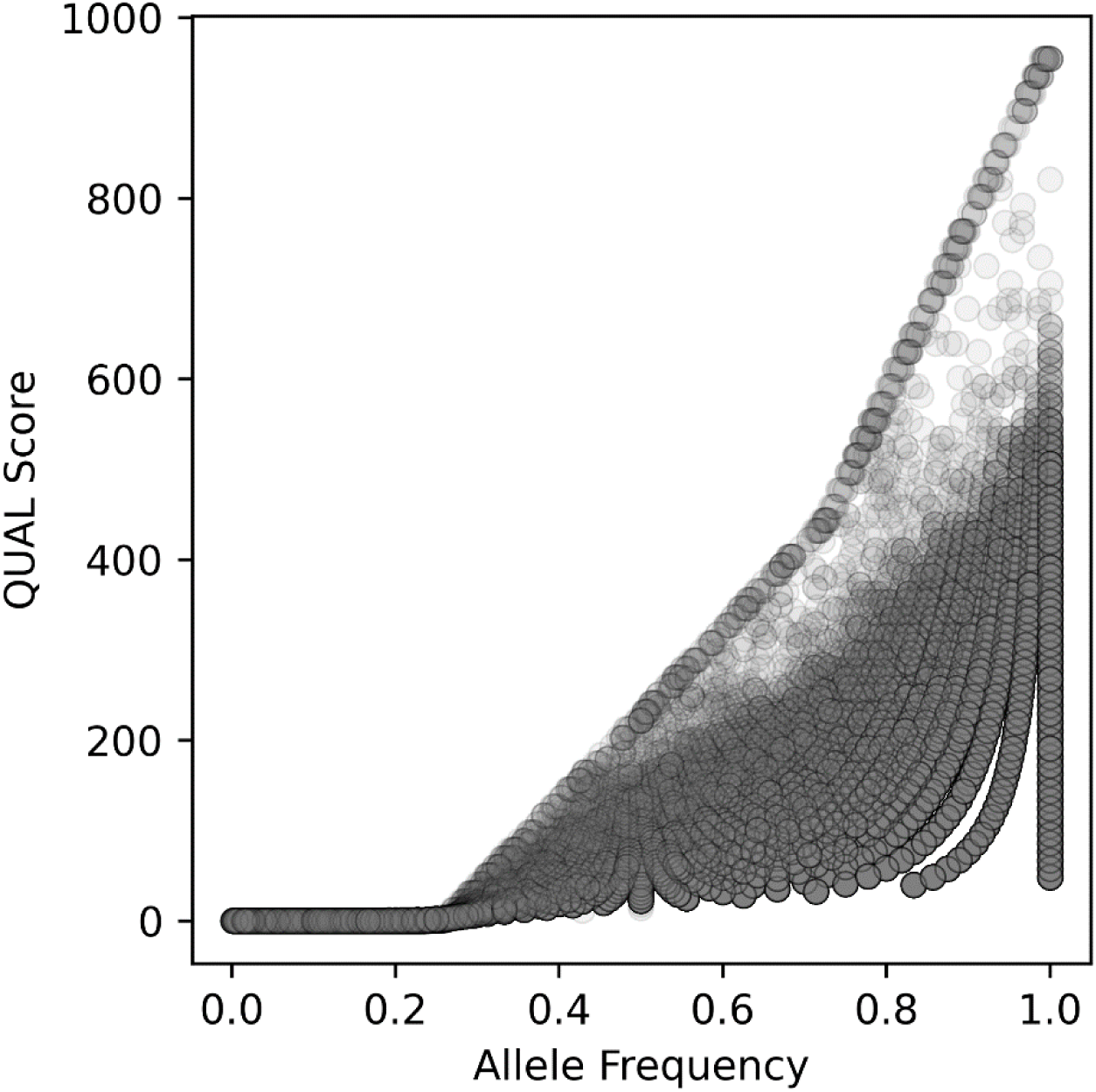
Commonly used metrics for filtering variants from individual samples are impacted by SV frequency within a population. Variants in a pooled sample can be present in a range of frequencies from 0-100% compared to single individual genotypes with expected to be homozygous (0% or 100%), or heterozygous (50%) (A) Quality (QUAL) scores use a logarithmic Phred scale to assess confidence in a variant call in the genome. QUAL scores are depressed (<10; indicating 90% confidence) in all variants until the allele frequency reaches a minimum of 27%. (B) Genotype quality (GQ) scores use a logarithmic Phred scale to assess confidence in a genotype call at a particular position in the genome.

**Supplemental Figure 2.**
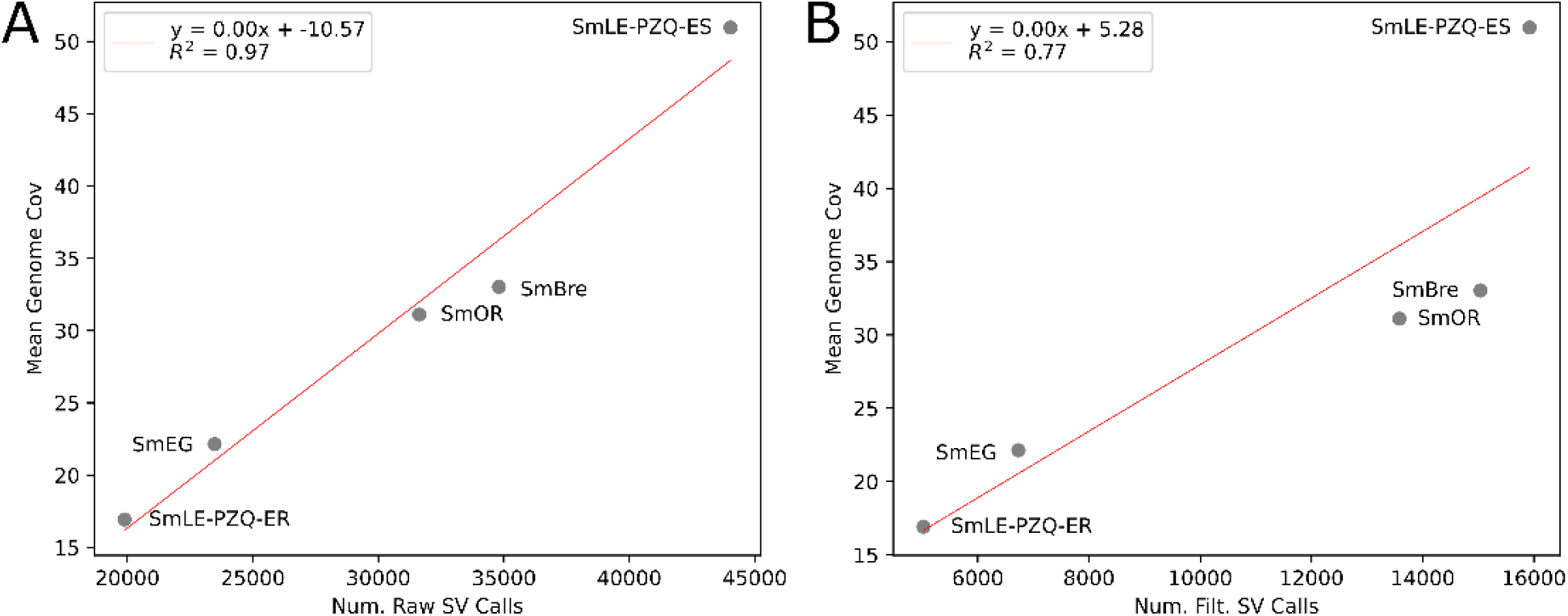
A positive relationship exists between the number of SVs identified de novo in each population and the amount of genome coverage in each of the five *S. mansoni* lab populations.

**Supplemental Figure 3.**
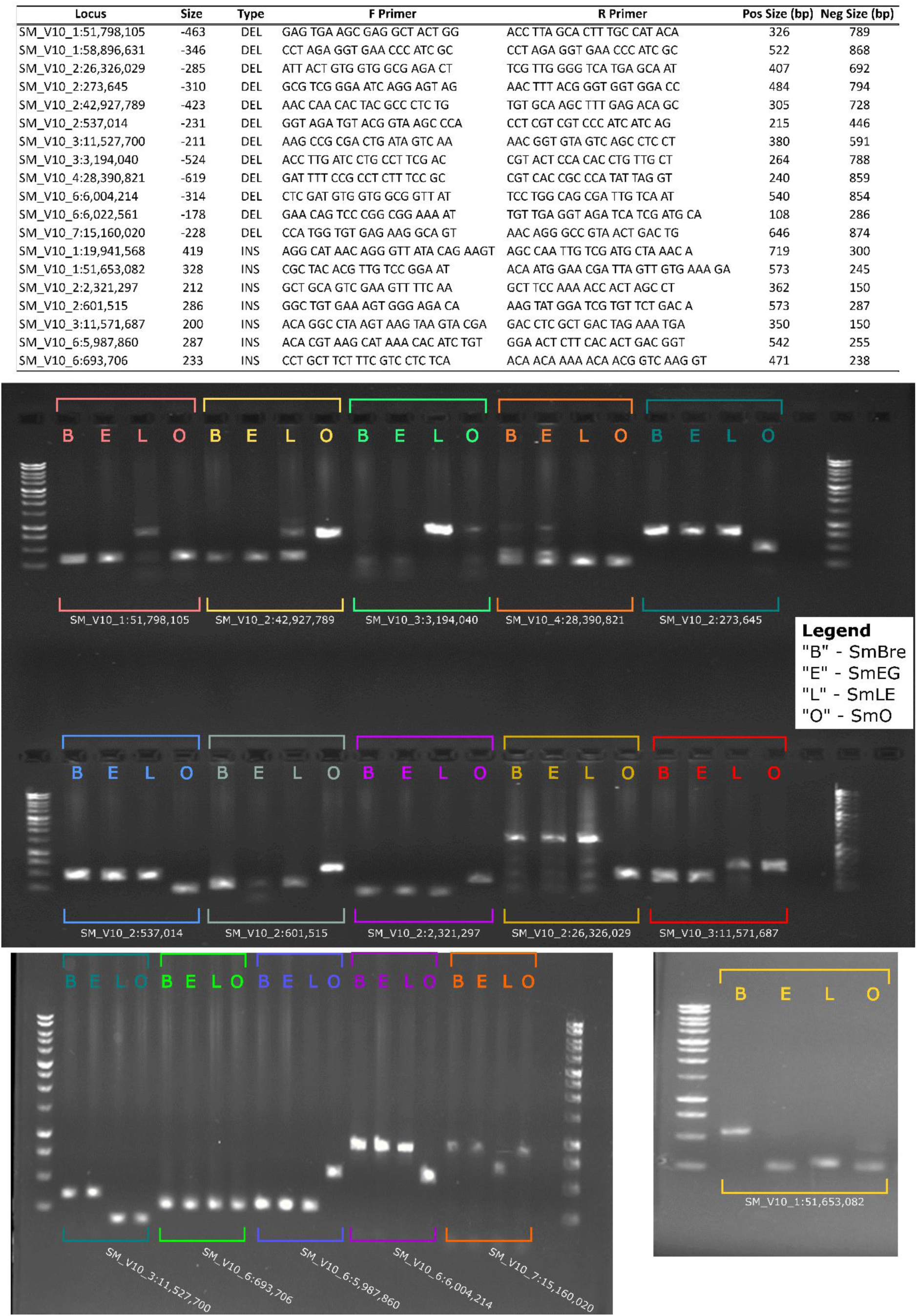
Structural variants (SV) were validated using PCR-based gel visualization. Nineteen of 25 loci were amplified (shown) and scored and the SV was scored as present or absent based on the amplicon size. Heterozygous individuals were identified by two bands.

